# Lipopolysaccharide-binding protein (LBP) can reverse the amyloid state of fibrin seen or induced in Parkinson’s disease: implications for its aetiology

**DOI:** 10.1101/124180

**Authors:** Etheresia Pretorius, Sthembile Mbotwe, Douglas B. Kell

**Affiliations:** Department of Physiological Sciences, Stellenbosch University, 7602, South Africa.; Department of Physiology, Faculty of Health Sciences, University of Pretoria, Arcadia 0007, South Africa.; School of Chemistry; The Manchester Institute of Biotechnology; Centre for Synthetic Biology of Fine and Speciality Chemicals, The University of Manchester, 131 Princess St, MANCHESTER M1 7DN, Lancs, UK

**Author notes:** **Corresponding authors: Etheresia Pretorius**, Department of Physiological Sciences, Stellenbosch University, Private Bag X1 MATIELAND, 7602, SOUTH AFRICA, **Douglas B. Kell** School of Chemistry and The Manchester Institute of Biotechnology, The University of Manchester, 131, Princess St, MANCHESTER M1 7DN, Lancs, UK.

**Keywords:** Thrombin, fibrin(ogen) clotting, LPS, LBP, thioflavin-T

## Abstract

The thrombin-induced polymerisation of fibrinogen to form fibrin is well established as a late stage of blood clotting. In recent work, we showed that the presence of tiny amounts of bacterial lipopolysaccharide (LPS) could cause these clots to adopt an amyloid form, that could be observed via scanning electron microscopy (SEM) or via the fluorescence of thioflavin-T. This could be prevented by the prior addition of lipopolysaccharide-binding protein (LBP). We had also observed by SEM this unusual clotting in the blood of patients with Parkinson’s disease (PD). We here show that this too can be prevented by LBP, thereby implicating such inflammatory microbial cell wall products in the aetiology of the disease. This may lead to novel treatment strategies in PD designed to target microbes and their products.

## Introduction

It is widely recognised that that many chronic, inflammatory diseases are accompanied by insoluble amyloid fibril formation (Chiti and Dobson, 2006; Herczenik and Gebbink, 2008; Rambaran and Serpell, 2008; Eisenberg and Jucker, 2012; Knowles et al., 2014; Tipping et al., 2015; Riek and Eisenberg, 2016). Thus, Parkinson’s (PD) is accompanied by amyloid forms of α-synuclein in the substantia nigra pars compacta (Uversky et al., 2001; Vilar et al., 2008; Olanow and Brundin, 2013; Kalia and Kalia, 2015; Kalia and Lang, 2015; Sampson et al., 2016). To this end, systems biology approaches (Antony et al., 2013; Funke et al., 2013; Jones et al., 2013; Fujita et al., 2014; Krishna et al., 2014) have made considerable headway in accounting for the known “Parkinson’s” genes in biochemical terms, with the additional recognition that iron dysregulation is also a major contributor to disease progression (Double et al., 2000; Levenson, 2003; Jones et al., 2007; Barnham and Bush, 2008; Gerlach et al., 2008; Perez et al., 2008; Altamura and Muckenthaler, 2009; Kell, 2009; Kell, 2010; Jameson, 2011; Oshiro et al., 2011; Weinreb et al., 2013; Kell and Pretorius, 2014).

It is rather less widely recognised that PD is also accompanied by major changes in the normal clotting of blood, i. e. it is a coagulopathy (Sato et al., 2003; Rosenbaum et al., 2013; Pretorius et al., 2014c; Infante et al., 2016).

When thrombin is added to the platelet poor plasma (PPP) of healthy controls, the fibres forming the subsequent clot appear like a plate of noodles or spaghetti in the scanning electron microscope (Campbell et al., 2010; Pretorius et al., 2011b; Weigandt et al., 2012; Bester et al., 2015; Kell and Pretorius, 2015b). However, we and others have observed that their diameter and morphology changes markedly in a variety of vascular and inflammatory diseases, typically producing ‘dense matter deposits’ (e.g. (Jörneskog et al., 1996; Dunn and Ariëns, 2004; Dunn et al., 2005; Dunn et al., 2006; Pieters et al., 2006; Alzahrani and Ajjan, 2010; Pretorius et al., 2011a; Pretorius et al., 2011b; Alzahrani et al., 2012; Pretorius and Kell, 2014; Pretorius et al., 2015)).

Thrombin removes two fibrinopeptides from fibrinogen, thereby allowing the fibrinogen to self-assemble into insoluble fibrin via a ‘knobs and holes’ mechanism (e.g. (Weisel, 2005; Wolberg, 2007; Cilia La Corte et al., 2011; Undas and Ariëns, 2011; Wolberg, 2012)). There are not otherwise considered to be any major changes in secondary structure (Weisel, 2005; Averett et al., 2008; Yermolenko et al., 2011; Protopopova et al., 2015). A final crosslinking step catalysed by Factor XIII (after it too has been activated by thrombin) (Dickneite et al., 2015) increases the stability of the clot.

A very specific feature of amyloid proteins is the formation of a cross-β-sheet structure, perpendicular to the fibres with a characteristic spacing (observable in X-ray reflections) of 4.7-4.8Å (e.g. (Maji et al., 2009; Eisenberg and Jucker, 2012; Tycko and Wickner, 2013; Riek and Eisenberg, 2016)). In contrast to normal structures, thioflavin T binds strongly to them, and fluoresces strongly at 480-520nm when excited at ~440 nm (e.g. (LeVine, 1999; Biancalana et al., 2009; Biancalana and Koide, 2010; Groenning, 2010; Sulatskaya et al., 2011; Freire et al., 2014)).

Although it can become so in the presence of a rare mutation in the fibrinogen a chain (Serpell et al., 2007; Picken, 2010; Stangou et al., 2010; Haidinger et al., 2013)), or by extreme mechanical stretching (Zhmurov et al., 2011; Litvinov et al., 2012; Zhmurov et al., 2012), fibrinogen is not considered to be amyloidogenic, nor is fibrin seen as an amyloid protein. However, following many observations in the SEM of anomalous blood clotting (e.g. (Pretorius et al., 2013b; Kell and Pretorius, 2014; Pretorius et al., 2014a; Pretorius and Kell, 2014; Bester et al., 2015; Kell and Pretorius, 2015b; Pretorius et al., 2015; Pretorius et al., 2016d)), we have recently established (Kell and Pretorius, 2016b; a; Pretorius et al., 2016c; Pretorius et al., 2016d) that this anomalous clotting is in fact amyloid in nature.

Dormant bacteria are widespread in nature (Kaprelyants et al., 1993; Domingue and Woody, 1997; Kell et al., 1998; Mattman, 2001; Domingue, 2010), and we have argued strongly for a role for dormant bacteria in the aetiology of such diseases (Kell et al., 2015; Kell and Pretorius, 2015a; Potgieter et al., 2015; Itzhaki et al., 2016; Kell and Kenny, 2016b; Kell and Pretorius, 2016c; Pretorius et al., 2016a; Pretorius et al., 2016b). In the recent analysis, amyloid formation occurred in the presence of tiny amounts of bacterial LPS, but was abolished when this was added together with a two-fold stoichiometric excess of human LBP (lipopolysaccharide binding protein) (Pretorius et al., 2016d). Recent work in mice lends strong support to the view that the gut microbiome can play a major role in the aetiology of PD (Sampson et al., 2016).

Iron is also capable of catalysing anomalous blood clotting (Pretorius et al., 2013b; Pretorius et al., 2014a), and there are strong indications for both iron dysregulation (Double et al., 2000; Kaur et al., 2003; Valko et al., 2005; Berg et al., 2008; Kell, 2008; Kell, 2010; Friedman and Galazka-Friedman, 2012; Funke et al., 2013; Hare et al., 2014; Kell and Pretorius, 2014; McDowall and Brown, 2016) and coagulopathies (Pretorius and Kell, 2014) in Parkinson’s disease. The purpose of the present paper was thus to study whether (the extent of) fibrin-type amyloid in blood varies between suitably matched controls and individuals with Parkinson’s disease. In addition, we wished to know whether the removal of any LPS using LBP affected this in any way. It became clear that the answers are in the affirmative in both cases.

## Materials and Methods

### Ethical statement

Ethical clearance was obtained from the Health Sciences Ethical Committee of the University of Pretoria and informed consent was obtained from each of the patients, as well as from controls (ethical number: 80/2013 and reapproved 2015). Exclusion criteria for the PD were conditions such as asthma, human immunodeficiency virus (HIV) or tuberculosis, and risk factors associated with metabolic syndrome, smoking, and (if female) being on contraceptive or hormone replacement treatment. Exclusion criteria for the healthy population were known inflammatory conditions such as asthma, human immunodeficiency virus (HIV) or tuberculosis, and risk factors associated with metabolic syndrome, smoking, and if female, being on contraceptive or hormone replacement treatment. This population did not take any anti-inflammatory medication. Whole blood of all participants was obtained in citrate tubes and platelet poor plasma (PPP) was used for confocal and SEM experiments. The methods were carried out in accordance with the approved guidelines. Blood was collected and methods were carried out in accordance with the relevant guidelines of the ethics committee. We adhered strictly to the Declaration of Helsinki.

### Sample population

In this study, 10 healthy, age-controlled individuals of whom 9 were spouses of some of the Parkinson’s disease (PD) individuals, and 26 individuals diagnosed with PD, were included. We also statistically compared results of healthy individuals from a previous study with Exclusion criteria for the healthy population were known inflammatory conditions such as asthma, human immunodeficiency virus (HIV) or tuberculosis, and risk factors associated with metabolic syndrome, smoking, and if female, being on contraceptive or hormone replacement treatment. The PD patients were diagnosed by a Neurologist and the Unified Parkinson’s Disease Rating Scale (UPDRS) was used in this diagnoses. On the day of blood collection, the Hoehn and Yahr scale was used by a clinician to rate the relative level of the PD disability (see Table 1 for the stages). Margaret M. Hoehn and Melvin D Yahr developed the Hoehn and Yahr scale to scale practically the severity of PD at the time of treatment, and thereby determine whether the medication or treatment that is used influences the rate of the progression of the disease (Hoehn and Yahr, 1967). Many studies thereafter have used this method in scaling the severity of movement disorders (Stocchi et al., 1997; Karlsen et al., 2000; Schrag et al., 2000a; b; Rodríguez-Violante et al., 2015).

**Table 1:** Relative level of disability and stage of Parkinson’s disease the Hoehn and Yahr scale.

| Stage | Symptoms |
| --- | --- |
| Stage 0 | No signs of disease |
| Stage 1 | Symptoms on one side only (unilateral) |
| Stage 1.5 | Symptoms unilateral and also involving the neck and spine |
| Stage 2 | Symptoms on both sides (bilateral) but no impairment of balance |
| Stage 2.5 | Mild bilateral symptoms with recovery when the 'pull' test is given (the doctor stands behind the person and asks them to maintain their balance when pulled backwards) |
| Stage 3 | Balance impairment. Mild to moderate disease. Physically independent |
| Stage 4 | Severe disability, but still able to walk or stand unassisted |
| Stage 5 | Needing a wheelchair or bedridden unless assisted |

### LPS-binding protein

A final LPS-binding protein (LBP) exposure concentration of 2 ng.L^-1^ LBP was used.

### Airyscan and scanning electron microscopy

PPP was prepared from whole blood collected in citrated tubes, from both healthy and PD individuals. For Airyscan preparation, we added Thioflavin T (ThT) at a final concentration of 5 µM to 200 µL of various prepared PPP samples and incubated it (protected from light) for one minute. This step was followed with the addition of thrombin, added in the ratio 1:2 to create extensive fibrin networks. A coverslip was placed over the prepared clot, and viewed immediately with a Zeiss LSM 510 META confocal microscope with a Plan-Apochromat 163 and 100x/1.4 Oil DIC objective. Excitation was at 488 nm and emitted light was measured at 505-550 nm. In addition, PPP from PD individuals were incubated with PBP (final concentration 2 ng. L^-1^) for 10 minutes, followed by ThT and clot preparation as for the healthy and naïve PD PPP. Clots were also prepared for SEM analysis, but after addition of thrombin, clots were washed, fixed in 4% formaldehyde and prepared for SEM according to known SEM preparation methods. Samples were viewed using a Zeiss cross beam electron microscope was used to study fibrin fibres.

### Statistical analysis

The non-parametric Mann–Whitney U test (between controls and PD samples) and the parametric T-test was performed (within samples) using the STATSDIRECT software.

## Results

Table 2 shows demographics for the healthy and the PD groups. The median of the Hoehn and Yahr scale for the PD individuals were 2.5 (±0.43); suggesting that mostly, individuals participating in this study had mild bilateral PD symptoms at date of blood collection. Airyscan results are shown in Figures 1 to 4, for the healthy and PD individuals.

**Figure 1:**
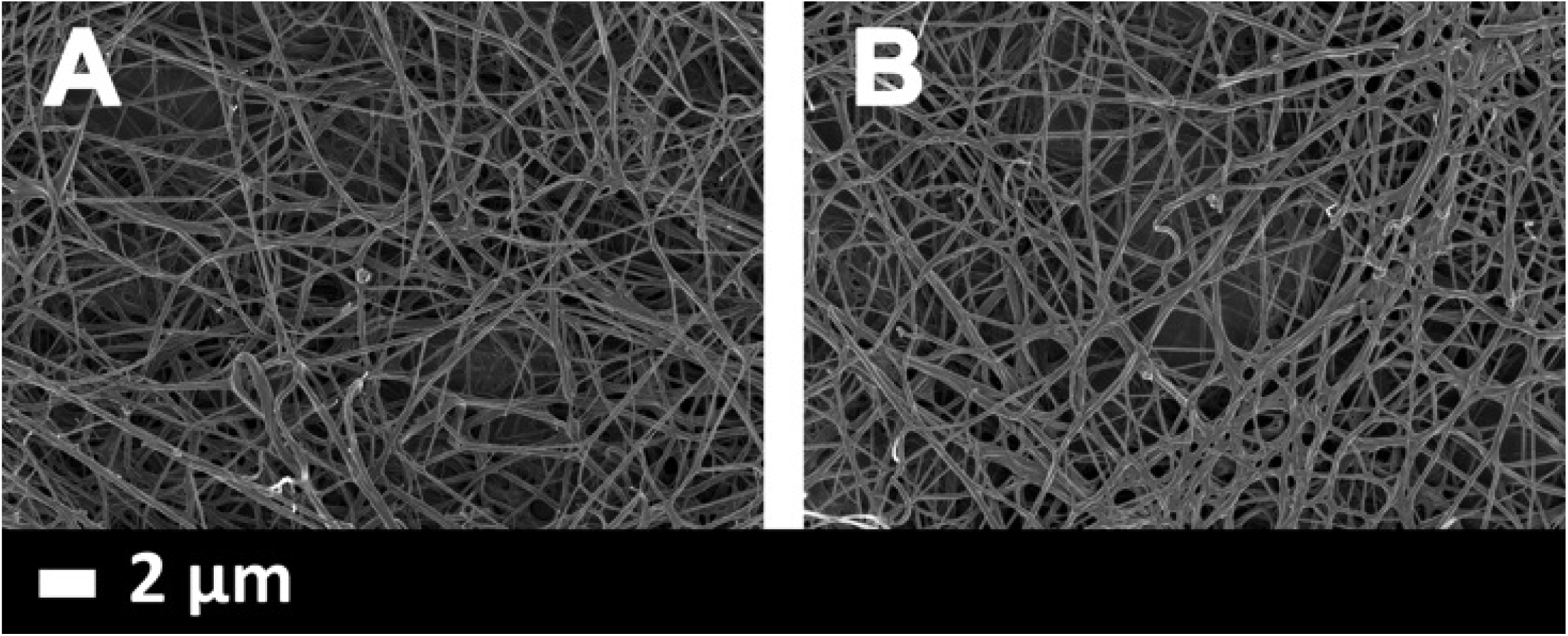
**A and B)** Clot structure from two healthy individuals. All clots were created by adding thrombin to PPP.

**Table 2:** Demographics for the healthy and the PD individuals.

| HEALTHY INDIVIDUALS (n=10) |  |  |
| --- | --- | --- |
|  | Gender | Age |
| Median; STD; % | 70% F; 30% M | All: 67.5 ($\pm 3$)<br>M: 67 ($\pm 2.5$)<br>F: 68 ($\pm 2.9$) |
| PARKINSON'S DISEASE INDIVIDUALS (Hoehn and Yahr scale 2.5 ($\pm 0.43$)) (n = 26) | | |
| Median; STD; % | 35% F; 65% M | All: 70 ($\pm 3.5$)<br>M: 70 ($\pm 3.9$)<br>F: 71 ($\pm 3.9$) |

### Scanning electron microscopy of PPP clots

Figure 1 shows how fibrin clots created from healthy PPP, looks like under a 10 000x machine magnification. Fibers have a spaghetti-like appearance, where individual fibres are visible. Fibrin clots created from PPP of PD individuals, all have a typical matted appearance, where individual fibres are entwined into a matted mass (see Figure 2), indicative of hypercoagulation. We recently suggested that this changed fibrin protein structure in inflammatory conditions like T2D, rheumatoid arthritis and others, are due to β-sheet-rich areas forming in the presence of both upregulated inflammatory markers, and particularly the presence or LPS (Pretorius et al., 2016d). We also showed that LPS added to healthy PPP created clots that showed hypercoagulation, and that the addition of LBP could reverse this pathological fibrin structure (Pretorius et al., 2016d). As referred to in the introduction, we also showed that the pathological fibrin structure of T2D could be reversed with the addition of LBP. Therefore, we suggested that LBP protects the fibres from LPS damage by binding to LPS and therefore that LBP could decrease β-sheet-rich areas in T2D plasma.

**Figure 2:**
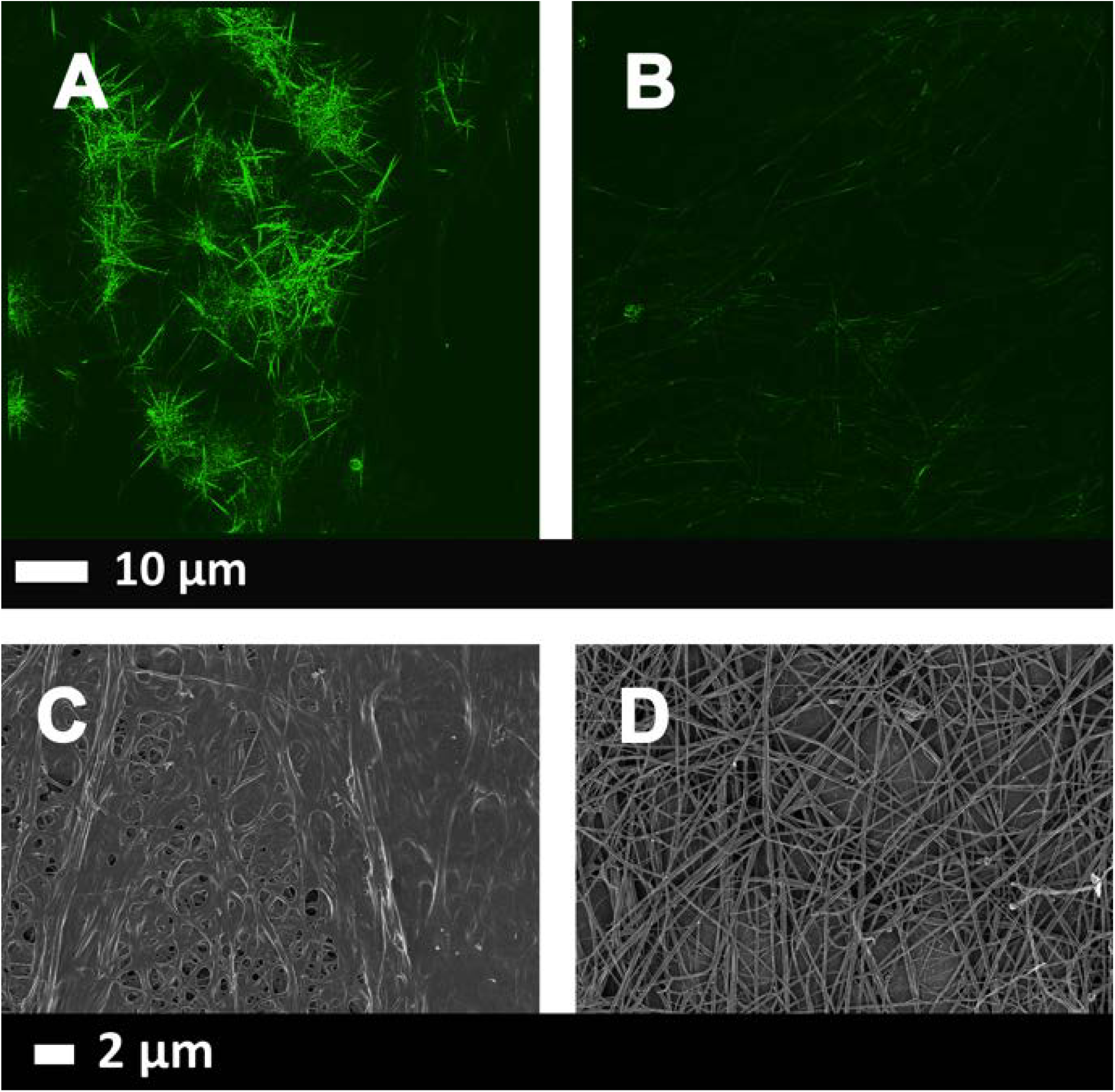
**A):** Airyscan micrographs of PPP with added thrombin to form extensive fibrin fibres from an individuals with Parkinson’s disease (PD); **B)** PPP from the same individual, but exposed to 2ng.L^-1^ LPS-binding protein followed by addition of thrombin. Thioflavin T (ThT) (5 µM) was added before thrombin. Micrographs were taken with a Zeiss LSM 510 META confocal microscope with a Plan-Apochromat 63×/1.4 Oil DIC objective. ***LBP dramatically reduced the fluorescence seen in samples from patients with PD.*** Gain settings were kept the same during all data capturing and not changed for statistical analysis, but brightness and contrast was slightly adjusted for figure preparation. **C)** SEM micrographs of PPP with added thrombin from the same individuals; **D)** PPP from same individual, but exposed to 2ng. L^-1^ LPS-binding protein followed by addition of thrombin.

Here we also added LBP to PPP from PD individuals. We could show that in all our PD samples, a structural revert to that similar to clots created from healthy PPP, could be obtained. All raw data, including extensive SOPs for SEM are stored on https://1drv.ms/f/s!AgoCOmY3bkKHmWY8VijKiQ-8_5RY and on EP’s researchgate profile,https://www.researchgate.net/profile/Etheresia_Pretorius.

### Airyscan super-resolution microscopy of clots created from PPP

Previously we have noted that in healthy PPP, in the presence of ThT, little to no fluorescence was present, with only occasional very small patches of fluorescence (Pretorius et al., 2016d)’(Kell and Pretorius, 2016a). We have also shown that when LPS had been added to healthy PPP, prior to thrombin, fluorescence was greatly enhanced, suggesting increased binding of ThT to β-sheet-rich areas on the fibrin(ogen) (Pretorius et al., 2016d)’(Kell and Pretorius, 2016a) From these results, we concluded that LPS binding causes the fibrinogen to polymerise into a form with a greatly increased amount of ß-sheet (in the presence of thrombin), reflecting amyloid formation. This causes a strong fluorescence observable (when excited ca 440 nm) in the presence of ThT (see e.g. (Biancalana et al., 2009; Biancalana and Koide, 2010)). In this paper, we also show β-sheet-rich areas in clots created from PPP of PD individuals (Figure 2).

Both Airyscan and SEM techniques are typically used only as qualitative methods. However, due to the increase fluorescence in clots prepared from PPP of PD, and also the more uniform and matted clot structure shown in SEM analysis of clots prepared from PD PPP, the variance between light and dark pixels are much less than seen in clots prepared from healthy PPP. We therefore propose using the coefficient of variation (CV) as our metric to quantify and discriminate between clots form healthy PPP and clots from PD PPP. We used ImageJ to calculate the mean and standard deviation of the intensity of the pixels in the images of the clot, using the histogram function, followed by the calculation of the coefficient of variation (i.e. SD/mean) of the intensity of the clot structure. Figure 3 shows boxplots of our SEM results and Figure 4A to D show examples of representative histograms of the 8-bit intensity for a typical SEM and confocal clot with and without LBP of a patient with PD. Within-sample analysis was done with the paired T-test and between-samples analysis was done using the Mann-Whitney test (See Table 3). We did not in this paper repeat Airyscan analysis with clots created with healthy PPP

**Figure 3:**
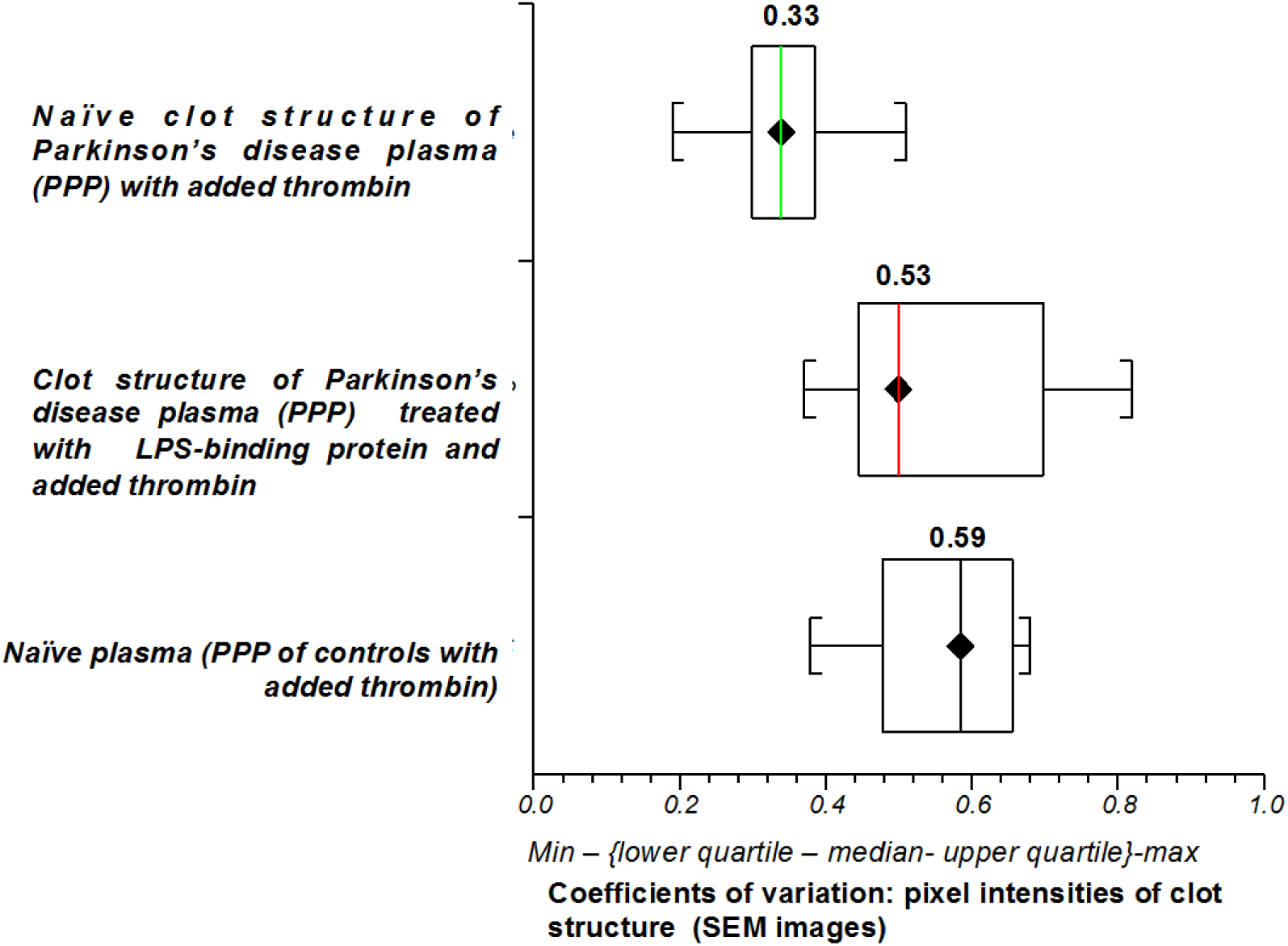
Boxplots of the distribution of the coefficients of variation in the pixel intensities of the SEM clot images from the different sample classes analysed (median coefficients of variation for each group is in box above plots).

**Figure 4:**
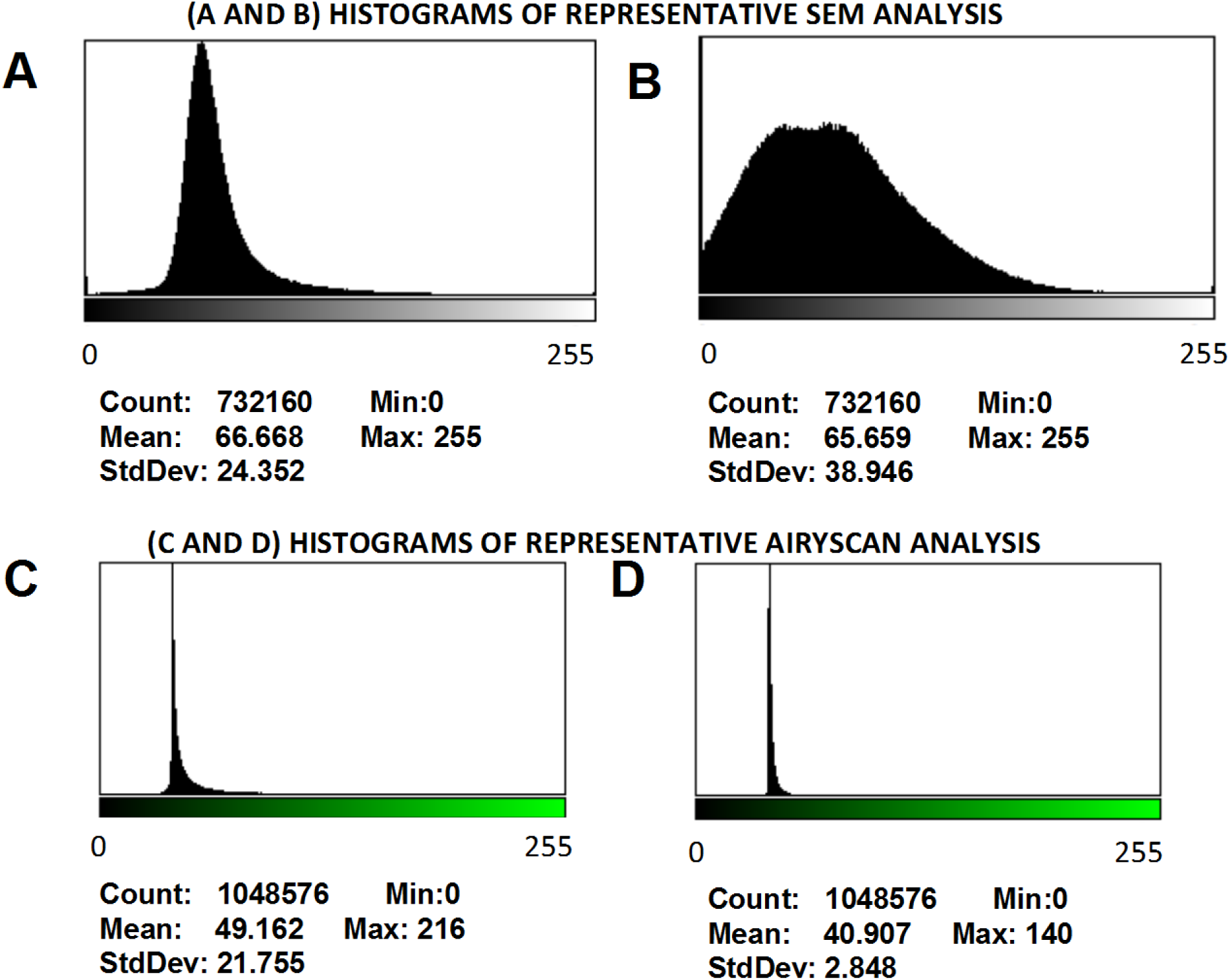
**A and B)** Representative histograms of the 8-bit intensity for a typical SEM clot from PPP of an individual with Parkinson’s disease and after addition of LBP. **C and D)** Representative histograms of the 8-bit intensity for a typical Airyscan clot from PPP of an individual with Parkinson’s disease and after addition of LBP.

**Table 3:**
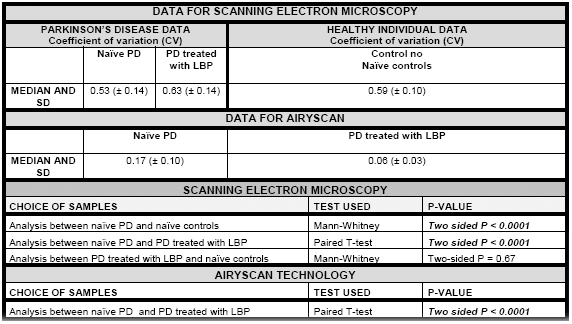
Data for Parkinson’s disease and healthy individuals showing the coefficients of variation (CV) of the intensity of the pixels in the clot images.

| DATA FOR SCANNING ELECTRON MICROSCOPY |  |  |  |
| --- | --- | --- | --- |
| PARKINSON'S DISEASE DATA<br>Coefficient of variation (CV) |  |  | HEALTHY INDIVIDUAL DATA<br>Coefficient of variation (CV) |
|  | Naïve PD | PD treated with LBP | Control no Naïve controls |
| MEDIAN AND SD | 0.53 (± 0.14) | 0.63 (± 0.14) | 0.59 (± 0.10) |
| DATA FOR AIRYSCAN |  |  |  |
|  | Naïve PD | PD treated with LBP |  |
| MEDIAN AND SD | 0.17 (± 0.10) | 0.06 (± 0.03) |  |
| SCANNING ELECTRON MICROSCOPY |  |  |  |
| CHOICE OF SAMPLES |  | TEST USED | P-VALUE |
| Analysis between naïve PD and naïve controls |  | Mann-Whitney | *Two sided P < 0.0001* |

|  |  |  |
| --- | --- | --- |
| Analysis between naïve PD and PD treated with LBP | Paired T-test | ***Two sided P < 0.0001*** |
| Analysis between PD treated with LBP and naïve controls | Mann-Whitney | Two-sided P = 0.67 |
| **AIRYSCAN TECHNOLOGY** |  |  |
| **CHOICE OF SAMPLES** | **TEST USED** | **P-VALUE** |
| Analysis between naïve PD and PD treated with LBP | Paired T-test | ***Two sided P < 0.0001*** |

## Discussion

Although we have observed anomalies in the kinds of fibrin fibres produced in the plasma of patients with various inflammatory diseases (e.g. (Pretorius et al., 2011b; Pretorius et al., 2011c; Pretorius et al., 2012; Pretorius et al., 2013b; Pretorius et al., 2014a; Pretorius et al., 2014b; Kell and Pretorius, 2015b; Pretorius et al., 2015; Kell and Pretorius, 2016a)), this is the first time that we have observed fibrin amyloid in Parkinson’s Disease, as assessed by thioflavin T staining and its sensitivity (and that of fibres observed in the SEM) to LBP. While fibrin and α-synuclein can coaggregate (Bhattacharjee and Bhattacharyya, 2014), it is especially notable that the thrombin-dependence and SEM fibre sizes tell us that the fibres we observe imply that they are essentially all made of fibrin.

Although α-synuclein fibre formation in the substantia nigra is characteristic of PD, it can also occur extracellularly, and its removal may be of therapeutic benefit (Kim et al., 2012; Park and Kim, 2013). The production may be driven by intestinal LPS (Kelly et al., 2014), while gut microbiota-derived short-chain fatty acids may also have a role (Sampson et al., 2016).

In terms of treatment (Kakkar and Dahiya, 2015; Kalia et al., 2015), we have previously discussed the potential role of iron chelators, both to stop the Fenton reaction (Kell, 2009)’(Kell, 2010) and to inhibit microbial proliferation (e.g. (Kell et al., 2015; Kell and Pretorius, 2015a; Potgieter et al., 2015; Kell and Kenny, 2016a)), and iron chelators can definitely also inhibit the formation of dense matted deposits (e.g. (Kell and Pretorius, 2014)’(Pretorius et al., 2013a; Nielsen and Pretorius, 2014; Pretorius et al., 2014a; Pretorius and Kell, 2014; Kell and Pretorius, 2015b)). It is now clear that treatment options worth exploring also include anticoagulants; as yet, however, the evidence for any effect of heparin is awaited, due to RCTs not having been done (Li et al., 2010)

## Concluding remarks

Overall, the remarkable reversal of amyloid fibrin formation by LBP addition to the plasma of Parkinson’s disease patients implies strongly that LPS is naturally pre-existing in said plasma. Although almost all the LPS is bound to plasma proteins under normal conditions (Kell and Pretorius, 2015a), including presumably to fibrinogen, but in concentrations that are consequently hard to determine (Kell and Pretorius, 2015a), it is known from the effects of adding them exogenously that LBP molecules can inhibit the LPS-induced formation of the amyloid form of fibrin when thrombin is present (Pretorius et al., 2016d). Consequently, the present work lends strong support to the idea (and evidence (Gabrielli et al., 2011; Tlaskalová-Hogenová et al., 2011)) that a dormant blood and tissue microbiome is at least part of the aetiology of Parkinson’s disease.

## Acknowledgements

We thank the Biotechnology and Biological Sciences Research Council (grant BB/L025752/1) as well as the National Research Foundation (NRF) of South Africa (91548: Competitive Program) and the Medical Research Council of South Africa (MRC) (Self-Initiated Research Program: A0X331 for supporting this collaboration. The authors thank Dr Prashilla Soma: Clinician. Some reagent funds were supplied as a philanthropic gift by an anonymous donor.

## Author contribution statement

EP is the study leader, analysed all samples, prepared all figures, co-wrote paper; SM: technical assistance with SEM preparation; DBK is the study co-leader, and co-wrote and edited the paper. All authors reviewed the manuscript.

